# Understanding associative false memories in aging using multivariate analyses

**DOI:** 10.1101/2021.07.26.453271

**Authors:** Nancy A. Dennis, Amy A. Overman, Catherine M. Carpenter, Courtney R. Gerver

## Abstract

Age-related declines in associative memory are ubiquitous, having been observed across a wide array of stimuli and experimental paradigms. Research further shows that such decreases in behavioral discriminability arise from increases in false memories for recombined lures. The current study examined the underlying neural basis of associative false memories by using representational similarity analyses to examine both age differences in the reactivation of encoded representations during retrieval and the neural overlap in the similarity of neural patterns underlying targets and related lures during retrieval. Behaviorally, we observed an age-related reduction in *d’*, which was shown to be driven by increased false alarms in the older adults. While we found no age difference in the relationship between patterns of neural activity underlying hits across memory phases (as measured by ERS), the similarity of neural patterns underlying targets and lures was affected by age. Specifically, while younger and older adults exhibited overall higher pattern similarity between hits and CRs compared to hits and FAs (as measured by RSA), this difference was reduced in older adults within occipital cortices. Additionally, greater Hit-FA representational similarity correlated with increases in associative FAs across occipital, frontal, and parietal cortices. Results suggest that while neural representations underlying targets may not differ across age, greater pattern similarity between the neural representation of targets and lures may reflect reduced distinctiveness of the information encoded in memory, such that old and new items are more difficult to discriminate, leading to more false alarms.

## Introduction

Associative memory is essential to daily life. It requires the ability to not only remember items and events of the past, but to remember the specific relationship between details of a given event. For example, it is important to remember both the faces and the names of new people you meet, as well as remembering which name is associated to each face. Misremembering one or more of these associative details, such as calling someone by the wrong name, can feel like a relatively minor memory error, but even these minor false memory errors can cause embarrassment and frustration. Associative memory success is most critical in cases where the consequences of errors are more serious, such as remembering medications and their accompanying dosages, a task that many older adults regularly encounter in their daily routines. Unfortunately, while associative false memories occur throughout the lifespan, they have been shown to be more frequent in older individuals (Chen & Naveh-Benjamin, 2012; Kilb & Naveh-Benjamin, 2007; Naveh-Benjamin, 2000; Naveh-Benjamin et al., 2004). Given that successfully navigating the world depends on making and retrieving accurate associations, it is important to understand how such associative false memories arise and what leads to increased error rates in older adults.

Age deficits in associative memory have been observed using a wide array of stimuli and experimental paradigms (e.g., Chalfonte & Johnson, 1996; Chen & Naveh-Benjamin, 2012; Kausler & Puckett, 1981a, 1981b; Kilb & Naveh-Benjamin, 2007; Naveh-Benjamin, 2000; Naveh-Benjamin et al., 2004; Naveh-Benjamin & Craik, 1995; Park et al., 1982; Park & Puglisi, 1985; Spaniol et al., 2006). The focus of this prior work has largely been on associative encoding and has examined how age-related deficits in binding contribute to memory errors (Earles et al., 2016; Nashiro & Mather, 2010; Naveh-Benjamin, 2000; Old & Naveh-Benjamin, 2008).

Specifically, behavioral and hippocampal evidence suggests that older adults exhibit a reduced ability to encode the relationship amongst discrete pieces of information, leading to associative information not being as well remembered as individual items (e.g., Bastin & Van der Linden, 2006; Chalfonte & Johnson, 1996; Dennis, Hayes, et al., 2008; Giovanello, Kensinger, Wong, & Schacter, 2009; Mitchell, Raye, Johnson, & Greene, 2006; Naveh-Benjamin, 2000; Sperling et al., 2003). While this deficit is largely conceptualized as an age-related reduction in associative hits (i.e., correctly recognizing, at retrieval, an associative pair previously seen at encoding), a closer look at the results of prior studies, our own included (Dennis, Hayes, et al., 2008; Overman, McCormick-Huhn, Dennis, Salerno, & Giglio, 2018), suggests that older adults’ associative memory deficit also arises from the endorsement of recombined lures (i.e., test pairs in which each individual stimulus had originally appeared as part of two different encoding pairs) during retrieval. This results in higher false alarm (FA) rates in older compared to younger adults (e.g., Castel & Craik, 2003; Healy, Light, & Chung, 2005; Naveh-Benjamin, Hussain, Guez, & Bar-On, 2003; Old & Naveh-Benjamin, 2008b; Overman & Stephens, 2013; Rhodes, Castel, & Jacoby, 2008).

False memories become increasingly pronounced in older adults when there is a high degree of perceptual overlap between studied targets and lures presented at retrieval (Koutstaal & Schacter, 1997; Norman & Schacter, 1997). This overlap leads to a reclassification of related lures as “remembered” at a similar rate as targets (Glanzer & Adams, 1985; Hockley, 2008). Univariate analyses assessing the neural basis of perceptual false memories in both young and older adults have shown there to be a high amount of overlap in the physical location of activated voxels across true and false memories. For example, both true and false memories activate regions within medial prefrontal cortex, angular gyrus, occipital cortices and the medial temporal lobe (e.g., Dennis, Bowman, & Peterson, 2014; McDermott, Gilmore, Nelson, Watson, & Ojemann, 2017; Schacter, Buckner, Koutstaal, Dale, & Rosen, 1997; Webb & Dennis, 2019). Such overlap in neural activation has been taken as an indication of the similarity in neural processes across the two trial types. But given the nature of univariate analyses, they have not been able to speak to an overlap in the nature of the neural processing across trial types.

One reason associative memory tasks are especially challenging to older adults is that the *items* in recombined lures are highly familiar to the participant, having been presented previously. Yet an individual must decide not simply that they have seen the items previously, but whether that specific *pair of items* was previously presented together. It has been suggested that higher rate of false associative memories in older adults is due to their failure to make use of successfully encoded perceptual details when confronted with lures at retrieval, particularly when those lures are novel pairs comprised of previously seen items (Bulevich & Thomas, 2012; Koutstaal, 2003; Mitchell, Ankudowich, Durbin, Greene, & Johnson, 2013).

Reflecting this difficulty, and consistent with recent work showing that reductions in neural distinctiveness underlie perceptual false memories (Bowman, Chamberlain, & Dennis, 2019; Lee, Samide, Richter, & Kuhl, 2019), we posit that older adults will show greater overlap in the similarity of neural patterns underlying hits and false alarms, compared to patterns underlying hits and correct rejections. That is, we predict that representational similarity will be greater for retrieval pairs that are behaviorally labeled as “old” compared to the overlap of targets and lures that are correctly differentiated. We also posit that this representational overlap between hits and false alarms will be related to false alarm rates in older adults. To this point, neural pattern similarity analysis in younger adults has shown that false memories are more prevalent when there is higher overlap in the representational similarity between targets and related lures in with cortical regions critical to memory processing (Chadwick et al., 2016; Wing et al., 2020). The above-mentioned multivariate studies examining false memories, as well as our own univariate meta-analysis of false memories (Kurkela & Dennis, 2016), suggests that perceptual processing regions, including inferior and middle occipital cortex, lateral temporal cortices, and frontoparietal cortices may be critical regions for investigating representational similarity differences between age groups, based on the importance of these regions in mediating false memories across a wide range of memory paradigms.

Given this proposed overlap between targets and lures, the transfer of information from encoding to retrieval is a key component to associative memory success, including the ability to discount new information containing some familiar aspects (i.e., lure pairs comprised of familiar items) by retrieving strong encoding traces of target information. Specifically, with regard to associative memory, not only do the individual items need to be encoded well and recapitulated at retrieval, but critically the relationship between them needs to be recapitulated in order to endorse the *target* pair (a re-presentation of target items in the exact same pairing) and reject a *lure* pair (a recombination of target items in a novel pairing). When the encoded association is not sufficiently reinstated at retrieval, a lure may be mistakenly endorsed, resulting in a false memory. The reactivation of this encoded representation can be examined by assessing the similarity of neural patterns between individual target trials across memory phases (i.e., encoding/retrieval) using encoding–retrieval similarity (ERS) analysis (e.g., Jonker, Dimsdale-Zucker, Ritchey, Clarke, & Ranganath, 2018; Kuhl, Rissman, Chun, & Wagner, 2011; Ritchey, Wing, LaBar, & Cabeza, 2013; Wing, Ritchey, & Cabeza, 2015). Previous work using ERS analysis within the domain of episodic memory has suggested that overlapping patterns of activation between encoding and retrieval correlates with both item and source memory success.

For example, research shows that ERS is capable of detecting the reinstatement of individual episodic memories in the hippocampus (Tompary, Duncan, & Davachi, 2016), as well as in other temporal cortex regions, the occipital lobe, and the frontoparietal cortex (Wing et al., 2015; Xiao et al., 2017). Additionally, within the domain of associative memory, work from our group suggests that one’s ability to generate similar neural patterns across memory phases differs with respect to the similarity of stimulus configuration across those memory phases (Gerver, Overman, Babu, Hultman, & Dennis, 2020). Examining age differences in ERS of targets helps us understand whether there are age differences in how the encoded representations of associative information are recapitulated at retrieval, and whether reductions in pattern similarity across memory phases contributes to age-related increases in false associative memories. We posit that, if older adults experience difficulty binding associative information during encoding, this would lead to deficits in recapitulating associative information during the retrieval process. To this end, we would expect to see age-related reductions in ERS within regions critical to perception and binding processes, including regions in the ventral visual cortex and hippocampus, with these reductions related to increases in false memory errors. Further, based on our previous work showing that ERS is modulated by the configural congruency between encoding and retrieval (Gerver et al., 2020), we posit that age-related changes in ERS may be greater for targets in which configural changes between encoding and retrieval reduce the familiarity of the associative pair across memory phases. Thus, the present study will investigate the neural basis of age-related increases in associative false memory by using representational similarity analysis to not only examine the overlap between hits and false alarms during retrieval, but also to examine whether reductions in target recapitulation is related to false memory errors.

## Methods

### Participants

Thirty-one right-handed young adults (the same sample as reported in Gerver et al. 2020) and thirty right-handed older adults were recruited from the Pennsylvania State University community and greater State College area, respectively. All participants provided written consent and received compensation for their participation. Participants were screened for history of psychiatric and neurological illness, head injury, stroke, learning disability, medication that affects cognitive and physiological function, and substance abuse. On the day of the study, all participants provided written informed consent for a protocol approved by the Pennsylvania State University Institutional Review Board. All participants were native English speakers with normal or corrected-to-normal vision. One older adult was excluded due to an inability of the analysis program to process their data and one older adult because they received the wrong version of the memory task. Three younger adults were removed from the study due to incomplete data (e.g., they stopped the task early), one because of excess movement in the scanner, and one because of misunderstanding the paradigm. Thus, the reported results are based on data from 26 young adult participants ranging from 18 to 25 years old (19 female; age: M = 20.5 years, SD = 1.98 years) and 28 older adult participants ranging from 63 to 83 years old (21 female; age: M= 71.04 years, SD = 6.02 years)^1^.

### Cognitive Assessments

All participants provided demographic information. In addition, older participants completed a short battery of cognitive assessments on the day of the study prior to being scanned to assess general cognitive function. The total time taken to complete all tests was less than 30 minutes and a break was given after tests and before scanning. The cognitive assessment battery included MMSE (*M*=29.52, *SD*=0.98), Digit Symbol Coding (*M*=12.43, *SD*=2.29), Symbol Copy (*M*=102.5, *SD*=23.76), Digit Span (*M*=12.48, *SD*=2.64), Letter-Number Sequencing (*M*=12.09, *SD*=3.11), WAIS-III Vocabulary (*M*=11.39, *SD*=1.68), and the Geriatric Depression Scale (*M*=0.86, *SD*=1.14) (Kurlowicz & Wallace, 1999; Wechsler, 1981; Yesavage et al., 1982). The stimuli and procedure were the same as that reported in Gerver et al. 2020, and are replicated here for completeness.

### Stimuli

Stimuli consisted of 170 color photographs of faces and 170 color photographs of scenes paired together. Face stimuli consisted of both male and females faces, each exhibiting a neutral expression, taken from the following online databases: the Color FERET Database (Phillips, Moon, Rizvi, & Rauss, 2000), adult face database from Dr. Denise Park’s laboratory (Minear & Park, 2004), the AR Face Database (Martinez & Benavente, 1998), and the FRI CVL face database (Solina, Peer, Batageli, Juvan, & Kovac, 2003). Scene stimuli consisted of outdoor and indoor scenes collected from an Internet image search. Using Adobe Photoshop CS2 Version 9.0.2 and Irfanview 4.0 (www.irfanview.com/), we edited face stimuli to a uniform size (320 × 240 pixels) and background (black), and scene stimuli were standardized to 576 × 432 pixels.

During encoding, half of the face–scene pairs were presented side by side; and the other half, with the face superimposed on top of the scene (see Figure 1). At retrieval, half of the pairs were presented in a manner congruent with their original contextual configuration; and the other half, in a contextual configuration that was incongruent with how it had been originally presented (e.g., at encoding was superimposed, at retrieval was side-by-side). Each encoding and retrieval block consisted of 34 total pairs. Within the 34 pairs per retrieval block, 10 were lures (five of which were side-by-side configuration and five of which were superimposed configuration) in which the face was rearranged with a different scene than was originally presented at encoding. During encoding, pairs were presented for 4 sec and for 4 sec at retrieval. A jittered ISI (2–8 sec) separated the presentation of each image. Each encoding and retrieval block lasted 4 min and 18 sec. A second version of this task was created to counterbalance the design such that the same stimuli were presented in the alternate configuration across versions (e.g., a face–scene pair presented side-by-side at encoding in Version A was presented as superimposed at encoding in Version B).

**Figure 1.**
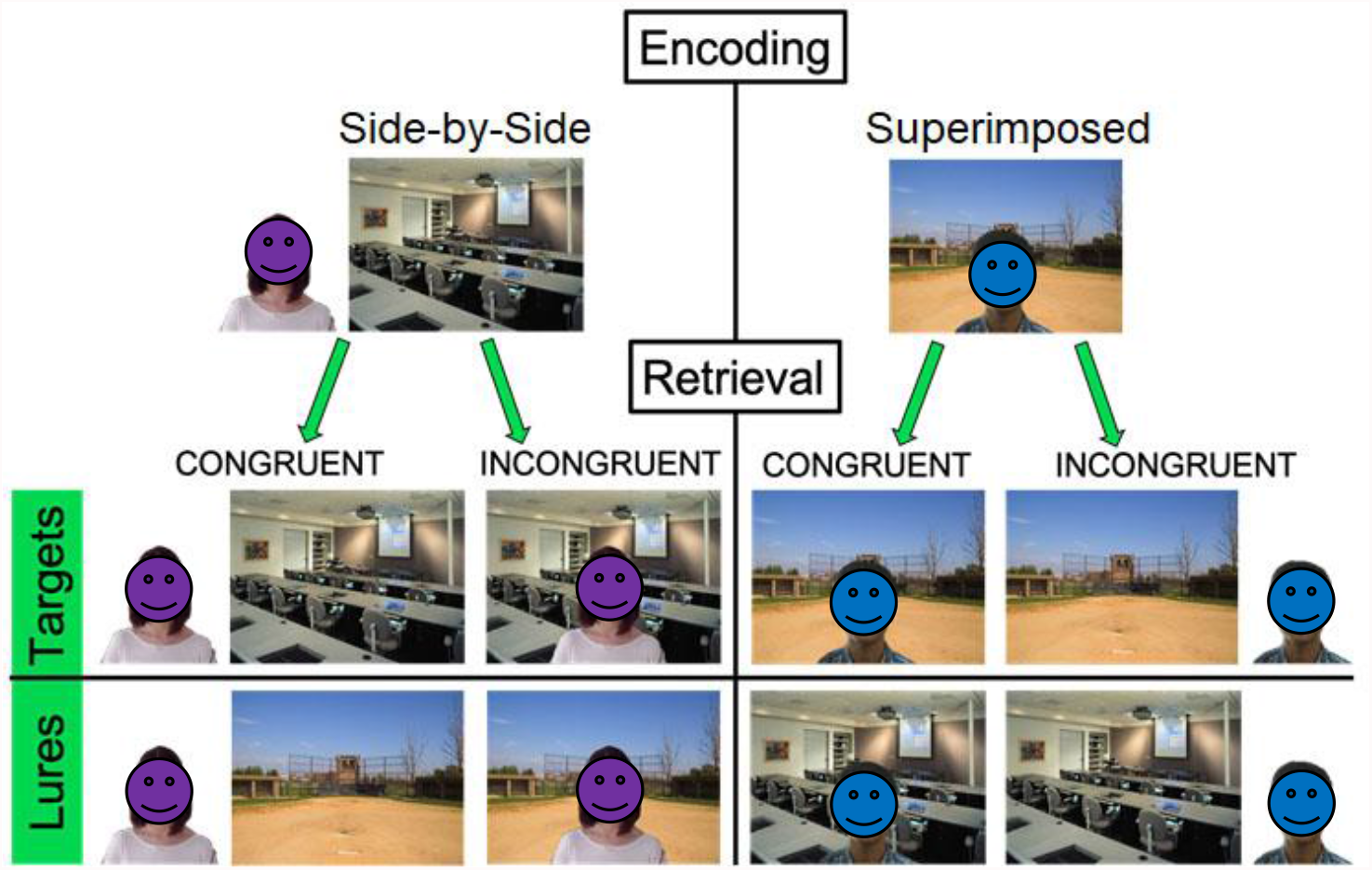
Task design. Examples of encoding and retrieval conditions for both congruent and incongruent targets and lures.

### Procedure

Before scanning, all participants were provided the full set of instructions for encoding and retrieval and completed a practice session of the full task. Participants were also asked to explain the instructions verbally before proceeding with the experiment to verify an accurate understanding of the task. They were encouraged to ask questions during this time. The scanning session began with a structural scan (magnetization prepared rapid gradient echo) that took approximately 7 min. During this time, the participants were asked to remain as still as possible. After the structural scan, participants completed five encoding and five retrieval blocks presented in an alternating order. Instruction screens, mirroring those used in the practice session, were presented before each block reiterating the verbal instructions participants received before entering the scanner. Presentation of all instruction screens was self-paced, meaning that the participants pressed “1” on the handheld button box to advance to the next screen when they read the instructions and were ready to begin the task.

After advancing past the instruction slides in encoding, the participant was presented with a series of face and scene pairings displayed on the screen in either a side-by-side or superimposed configuration. Each pair was presented for 4 sec, during which time the participant responded to the question “How welcoming are the scene and face together?” Text of the question was presented below each pair by using a rating scale from 1 to 4 (1= not at all, 4=very) and making a key-press on their handheld button box (See Figure 1a). The instructions emphasized that the participants should choose the rating based on how welcoming the face and scene pairing was collectively in order to facilitate encoding of both the face and the scene, rather than only one or the other, or them separately. This question also helped ensure that participants paid attention to the scene, even when it was configured behind the face in the superimposed condition because we did not want the scene to be incidentally encoded while the face was intentionally encoded. Across versions, faces and scenes were counterbalanced for their inclusions in either pair configuration. No differences across versions were noted, and all analyses were collapsed across versions.

Each encoding block was followed by a retrieval block. Similar to encoding, each face– scene pair at retrieval was presented for 4 sec. During this time, participants were asked to respond to the question: “Please identify the pairings that have been presented previously.” Displayed below the question were the following choices: 1 = remember, 2 = know, and 3 = new. A remember–know–new design was chosen to isolate recollection-related activity, associated with “remember” responses, from that of familiarity, associated with “know” responses. This distinction has shown to be critical when assessing memory-related activity particularly within the MTL (Yonelinas, 2002; Yonelinas et al., 2005, 2007). Participants were instructed to make their memory judgments based on the co-occurrence of the face and scene and not to base their judgments on the configuration of the display. Specifically, participants were instructed to select “Remember” if they were able to retrieve details of the face and scene appearing together at study, select “Know” if they thought the face and scene appeared together previously but they could not remember specific details about the original appearance, and select “New” if they believe that the face–scene pair did not appear together previously. Thus, whether a retrieval configuration was congruent or incongruent with how it had been presented at encoding, the Remember/Know/New labels applied equally to both types of configurations, with a participant’s response simply dependent on the vividness of their memory. Similar to the encoding phase, all responses were made using the button. With respect to retrieval configurations, half of the trials were congruent with their encoding configuration and half were incongruent (such that a side-by-side trial at encoding was presented as a superimposed trial at retrieval) (See Figure 1).

### Image Acquisition

Structural and functional images were acquired using a Siemens 3-T scanner equipped with a 12-channel head coil, parallel to the AC–PC plane. Structural images were acquired with a 1650-msec repetition time, a 2.03-msec echo time, a 256-mm field of view, a 2562 matrix, 160 axial slices, and a 1.0-mm slice thickness for each participant. Echo-planar functional images were acquired using a descending acquisition, a 2500-msec repetition time, a 25-msec echo time, a 240-mm field of view, a 802 matrix, a 90° flip angle, and 42 axial slices with a 3.0-mm slice thickness resulting in 3.0-mm isotropic voxels.

### Image Processing

Raw anatomical and functional brain images were first skull stripped using the Brain Extraction Tool (Smith, 2002) in the FMRIB Software Library (FSL) Version 5.0.10 (www.fmrib.ox.ac.uk/fsl). FSL’s MCFLIRT function (Jenkinson et al., 2002, 2012) was then applied for realignment and motion correction within each functional run. All volumes were aligned to the middle volume of the middle run of encoding. The realigned functional images were then processed by FSL’s fMRI Expert Analysis Tool (Woolrich et al., 2001), where they were high-passed filtered and spatially smoothed at 6-mm FWHM. These data were then pre-whitened to account for temporal autocorrelations within voxels. Finally, the structural data underwent nonlinear transformation into the standardized Montreal Neurological Institute (MNI) space by using the warping function in FSL’s FNIRT (Andersson et al., 2007). For multivariate analyses, the raw data underwent the exact same steps as above, absent smoothing.

### Behavioral Analyses

All behavioral analyses were conducted in Rstudio using base R functions and rstatix (www.rstudio.com; Kassabara, 2020). Because this study aimed to examine false associative memory, behavioral and neuroimaging analyses were focused on accurately and inaccurately recollected memory decisions, in the form of d’ (calculated in Excel as norm.s.inv(hits) – norm.s.inv(FAs)). To test for age differences in memory, three 2 (Between Age Group: Old, Young) × 2 (Within Encoding Configuration: side-by-side or superimposed) x 2 (Within Retrieval Congruency: congruent or incongruent) mixed model ANOVAs were performed. To examine significant main effects and interactions, paired-sample t-tests were run, with Bonferroni correction.

### Multivariate Pattern Similarity Analyses

To estimate neural activity associated with individual trials, separate GLMs were estimated in SPM12 defining one regressor for each trial at encoding and retrieval (170 in total for each respective memory phase) (Mumford, Davis, & Poldrack, 2014). An additional six nuisance regressors were included in each run corresponding to motion. Whole-brain beta parameter maps were generated for each trial at retrieval for each participant. In any given parameter map, the value in each voxel represents the regression coefficient for that trial’s regressor in a multiple regression containing all other trials in the run and the motion parameters. These beta parameter maps were next concatenated across runs and submitted to the CoSMoMVPA toolbox (Oosterhof et al., 2016) for similarity analyses (representational similarity analysis: RSA and encoding-retrieval similarity analysis: ERS).

### ROIs

Based on our focus on using ERS and RSA to better understand age-related increases in false memories, all analyses were computed in ROIs previously identified as critical to false memories. Specifically, we drew on regions identified both in past multivariate studies of false memories (Bowman et al., 2019; Lee et al., 2019; Wing et al., 2020) and our meta-analysis of neural activation underlying false memories (Kurkela & Dennis, 2016). ROIs used in the current study included the medial superior PFC (Medial SFG; *N*_voxels_ = 9,644), angular gyrus (AG; *N*_voxels_ = 801), inferior (IOC; *N*_voxels_ = 1,930) and middle occipital cortex (MOC; *N*_voxels_ = 5,368), and medial temporal gyrus (MTG; *N*_voxels_ = 10,013). We also included a subregion of the MTL, the bilateral hippocampus (HC; *N*_voxels_ = 1,878), that has been shown to be critical to associative memory and binding. While the general ROIs were drawn from this past work, specific ROIs in the current investigation were defined using AAL PickAtlas in SPM12 using anatomically defined boundaries identified by the anatomical labeling. The medial superior PFC region was created in the AAL PickAtlas based upon coordinates and activation identified in the Kurkela & Dennis meta-analysis.

### ERS analysis

Encoding-retrieval similarity analysis (Kriegeskorte, Mur, & Bandettini, 2008; Ritchey et al., 2013) was conducted to examine whether there was an age deficit in how encoded information was re-represented at retrieval and whether this influenced false memory performance, we directly compared neural patterns of activation between encoding and retrieval. On the basis of our previous work indicating that encoding configuration (i.e., side-by-side, superimposed) significantly influences patterns of neural activity during encoding (Dennis et al., 2019) and configural congruency between encoding and retrieval was critical to ERS (Gerver et al., 2020), we separated all targets into four conditions of interest: side-by-side congruent targets, superimposed congruent targets, side-by-side incongruent targets, and superimposed incongruent targets. To compute pattern similarity, we used the beta estimates extracted from the single trial models for encoding and retrieval described above. Activation for each individual trial for a given condition at encoding was Fisher’s z transformed and correlated with every trial of the same type at retrieval (e.g., targets at encoding that were presented as a side-by-side configuration that were congruently re-presented as side-by-side at retrieval). This resulted in similarity scores, as operationalized by Pearson’s *r* correlation values, for each trial. The correlations were then averaged within condition for each participant.

From these values, group averages were computed and age differences were assessed using a mixed factorial ANOVA, using the rstatix package in R (Kassambara, 2021), with age (young, old) as the between subjects variable and ROI (IOC, MOC, AG, MTG, medial SFG, hippocampus) and condition (side-by-side congruent, superimposed congruent, side-by-side incongruent and superimposed incongruent) as within subjects variables. To examine significant main effects and interactions, paired-sample t-tests were run and Bonferroni corrected using the emmeans R package (Lenth, 2021). Noted below in results, the ANOVA revealed no age differences in ERS. Using the BayesFactor package in R (Morey, Rouder, Jamil, & Morey, 2015) we provide Bayes factors that indicate how strong the evidence is for the null hypothesis (no age difference) relative to the alternative hypothesis (age difference) for ERS, collapsed across condition and ROI using an approach similar to Bowman, Ashby, and Zeithamova (2021). We did the same for our follow up ERS ANOVAs with ROIs when examining the absence of age effects in the main analyses.

### RSA

A representational similarity analysis (Kriegeskorte et al., 2008) was conducted to examine overlap in the representation of stimuli associated with different behavioral outcomes. Specifically, given our interest in understanding whether older adults’ increase in false memories was related to heightened pattern similarity across hits and FAs, we examined the similarity of patterns of neural activity between targets and related lures as a function of behavior. Specifically, we correlated neural pattern similarity between Hits and FAs (representing pattern similarity between different types of “old” responses; i.e., veridical and false memories) and between Hits and CRs (representing pattern similarity between different types of veridical memories) within each ROI described above. Given trial counts across each condition and the fact that false memories were not affected by contextual congruency, we collapsed across this factor in the current analysis. To compute pattern similarity, we used the beta estimates extracted from the retrieval single trial model described above to run two separate Pearson correlations between Hits and FAs and Hits and CRs. The resulting correlation matrices were Fisher’s z transformed then averaged to get a single correlation value per trial pair (Hits and FAs and Hits and CRs; see Trelle et al., 2019 for a similar methodological approach) for each participant.

From these values group averages were computed and age differences assessed using a mixed factorial ANOVA, using the rstatix package in R (Kassambara, 2021), with age (young, old) as the between subjects variable and ROI (IOC, MOC, AG, MTG, medial SFG, hippocampus) and trial pair (Hit and CR RSA, Hit and FA RSA) as within subjects variables. To examine significant main effects and interactions, paired-sample t-tests were run and Bonferroni corrected.

## Results

### Behavioral results

For the *d’* measures, the ANOVA revealed a significant main effect of age group, *F*(1,52)=16.95, *p*<.001, η^2^ = 0.25, such that young adults (*M*=1.40, 95% C.I. [1.32, 1.49]) had better discrimination than older adults (*M*= 0.68, 95% C.I. [0.60, 0.75]).The main effect of encoding configuration was also significant, *F*(1,52)=5.80, *p*=.020, η^2^ = 0.10, such that superimposed trials (*M*=1.12, 95% C.I. [1.03, 1.20]) had better discrimination between targets and lures than side-by-side trials (*M*=0.93, 95% C.I. [0.84, 1.02]). The main effect of retrieval congruency and the two and three-way interactions were not significant (all *F*’s < 3.14, all *p*<u>’s</u> > .08).

Breaking the *d’* measure down by hit and FA rates we found that the hit ANOVA revealed a significant main effect of encoding configuration, *F*(1,52)=6.10, *p*=.017, η^2^ =0.11, such that superimposed configurations (*M*=0.84, 95% C.I. [0.83, 0.85]) had higher hits than side-by-side (*M*=0.82, 95% C.I. [0.81, 0.83]). Additionally, there was a significant main effect of retrieval configuration congruency, *F*(1,52)=53.83, *p*<.001, η^2^ =0.51, such that congruent trials (*M*=0.87, 95% C.I. [0.86, 0.88]) showed higher hit rates than incongruent trials (*M*=0.79, 95% C.I. [0.77, 0.80]). The main effect of age group and the two and three-way interactions were not significant (all *F*’s<1.26, all *p*’s>.27). With regard to FA rates, the ANOVA revealed a significant main effect of age group, *F*(1,52)=19.27, *p*<.001, η^2^ = 0.27, such that older adults (*M*=0.63, 95% C.I. [0.60, 0.65]) had higher false alarms than younger adults (*M*= 0.39, 95% C.I. [0.37, 0.41]). The main effect of retrieval configural congruency was also significant, *F*(1,52)=22.18, *p*<.001, η^2^ =0.30, such that congruent trials (*M*=0.55, 95% C.I. [0.52, 0.57]) showed higher false alarms than incongruent trials (*M*=0.48, 95% C.I. [0.45, 0.50]). The main effect of encoding congruency and the two and three-way interactions were not significant (all *F*’s < 3.67, all *p*’s >.06). [See Table 1 for means].

**Table 1.**
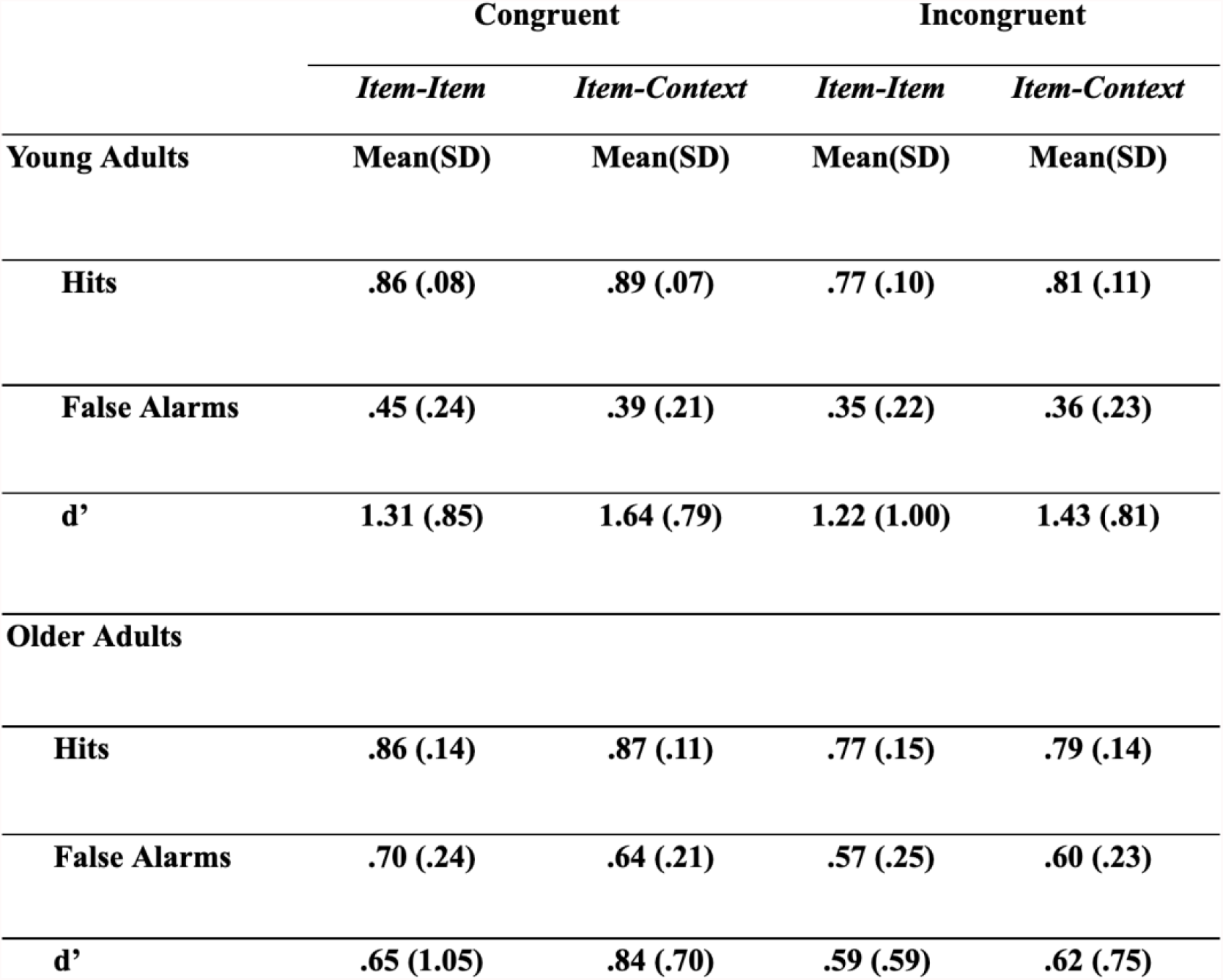
Behavioral performance broken down by condition and age group. Mean and standard deviation reported.

### ERS

The main objective in analyses of ERS was to examine age differences across conditions. Thus, the post-hoc analyses based on the ERS ANOVA were focused on meaningful comparisons with respect to those factors. Critically, the ERS ANOVA revealed both a significant main effects of condition and ROI and a significant 3-way interaction between condition, age, and ROI. With respect to the main effect of condition, *F*(3,153)=28.42, *p*<.001, both congruent side-by-side *(M*=0.010, *SD*=0.019) and congruent superimposed *(M*=0.011, *SD*=0.020) ERS metrics were greater than either of the incongruent ERS metrics (side-by-side *M*=0.0077, *SD*=0.021; superimposed *M*=0.0081, *SD*=0.021). Specifically, superimposed congruent had greater ERS than superimposed incongruent, *t*(52)=6.24, *p*<.001, and side-by-side incongruent, *t*(52)=6.99, *p*<.001. Additionally, side-by-side congruent showed greater ERS than superimposed incongruent, *t*(52)=5.05, *p*<.001, and side-by-side incongruent, *t*(52)=5.83, *p*<.001. There was also a significant main effect of ROI, *F*(2.13, 108.73)=21.20, *p*<.001. There was also a significant two-way interaction between Condition and ROI, *F*(4.89, 249.32), *p*<.001 and a three-way interaction between age, condition and ROI was significant *F*(4.89, 249.32)=2.46, *p*=.035. The main effect of age *F*(1,51)=0.75, *p*=.39, the two-way interaction between age group and condition *F*(3, 153)=1.10, *p*=0.35, interaction between age group and ROI *F*(2.13, 108.73)=0.85, *p*=.44, was not significant. Consistent with the findings from the ANOVA, a Bayesian analysis of age differences across ERS collapsed across condition and ROI yields modest evidence in favor of the null hypothesis (JZS BF_01_ = 1.11).

To examine whether the three-way interaction between condition and ROI and age was meaningful, we conducted a repeated measures ANOVA within each ROI. There was a significant main effect of condition within the IOC, *F*(3,156)=17.75, *p*<.001 and MOC, *F*(3,156)=43.06, *p*<.001. Follow-up paired-sample t-tests revealed that, within the IOC, ERS significantly differed between the side-by-side congruent (*M*=0.002, 95% C.I. [0.002, 0.003]) and side-by-side incongruent targets (*M*=-0.002, 95% C.I. [-0.003, −0.001]), *t*(52)= 4.59, *p*<.001, as well as superimposed incongruent targets (*M*=-0.002, 95% C.I. [-0.002, −0.001]), *t*(52)= 4.07, *p*<.001. Additionally, ERS significantly differed between the superimposed congruent targets (*M* =0.004, 95% C.I. [0.003, 0.005]) and the superimposed incongruent targets (*M*=-0.002, 95% C.I. [-0.002, −0.001]), *t*(52)= 5.08, *p*<.001 as well as side-by-side incongruent targets (*M*=-0.002, 95% C.I. [-0.003, −0.001]), *t*(52)= 5.33, *p*<.001. See Figure 2a. A similar pattern was found in the MOC where ERS significantly differed between side-by-side congruent targets (*M*=0.005, 95% C.I. [0.005, 0.006]) and side-by-side incongruent targets (*M*=-0.004,) 95% C.I. [-0.005, − 0.003], *t*(52)= 7.43, *p*<.001, superimposed congruent targets (*M*=0.008, 95% C.I. [0.007, 0.009]), *t*(52)= −3.01, *p*<.001, and superimposed incongruent targets (*M*=-0.004, 95% C.I. [-0.005, −0.002]), *t*(52)= 6.58, *p*<.001. Additionally, ERS significantly differed between superimposed congruent targets (*M*=0.008, 95% C.I. [0.007, 0.009]) and superimposed incongruent targets (*M*=-0.004, 95% C.I. [-0.005, −0.002]), *t*(52)= 6.73, *p*<.001 and side-by-side incongruent targets (*M*=-0.004, 95% C.I. [-0.005, −0.003]), *t*(52)= 7.78, *p*<.001. See Figure 2a. While neither ANOVA exhibited a main effect of age, a Bayesian analysis of age differences across ERS within the IOC collapsed across condition yields moderate evidence in favor of the alternative hypothesis (age differences) (JZS BF_10_ = 8.96), while a Bayesian analysis of age differences across ERS within the MOC collapsed across condition yields modest evidence in favor of the null hypothesis (no age differences) (JZS BF_01_ = 1.57).

**Figure 2.**
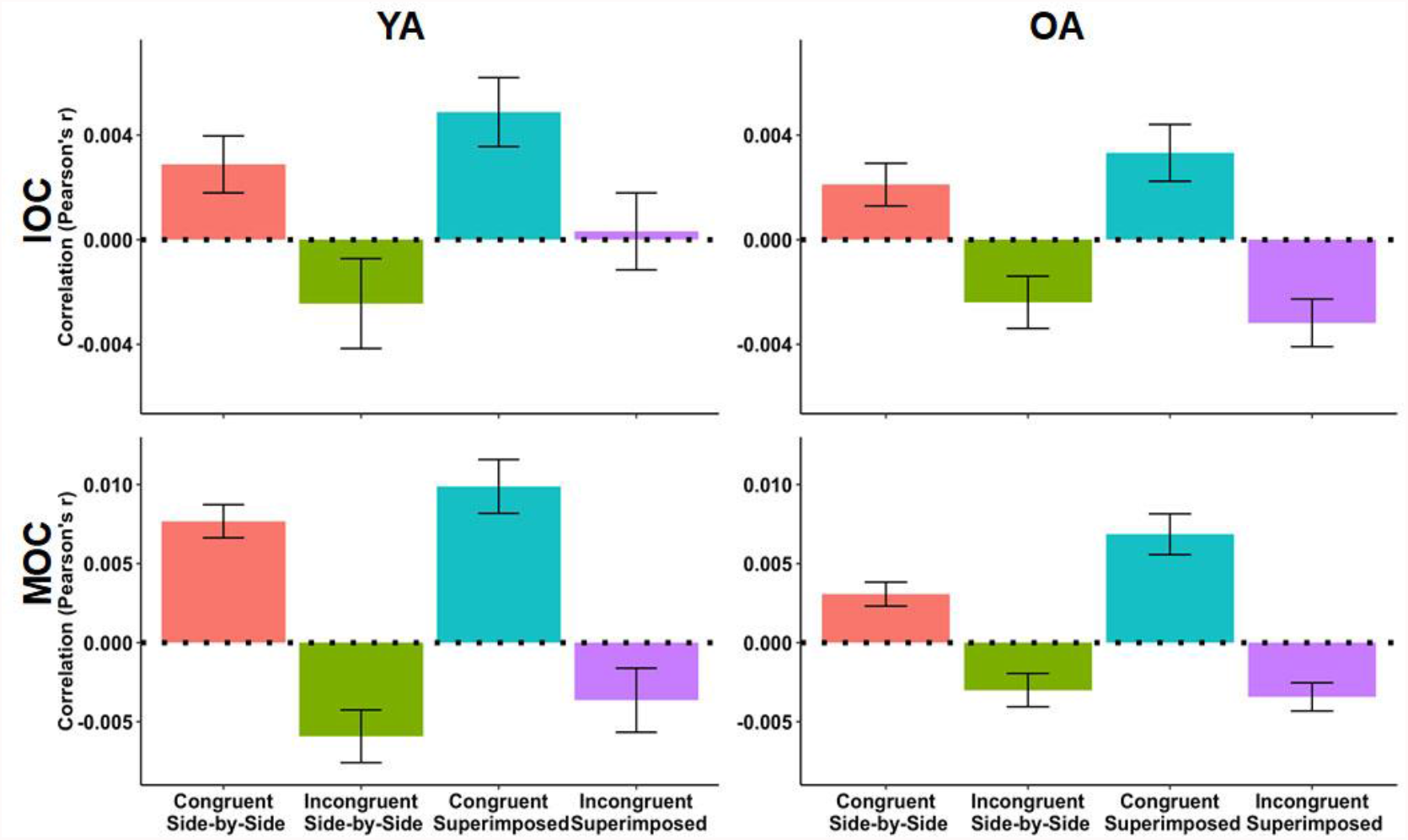
ERS in the Inferior Occipital Cortex (IOC) and Middle Occipital Cortex (MOC), broken down by the four target conditions in the younger and older adults. Error bars represent standard error of the mean.

There was no main effect of age in any ROI, suggesting that neural pattern similarity of all four contextual and configural conditions across encoding and retrieval yielded comparable ERS between young and older adults across all *a priori* regions of interest (all *p*’s > .05). This also suggests that the main effect of age within the 3-way interaction resulted from differences across condition and ROI which would not be interpretable within the current analysis/design. See Supplemental Table A for ERS values reported for each age group for each ROI.] Finally, we found no correlation between ERS metrics and memory performance in either age group. [See Figure 2].

### RSA

Similar to the ERS analysis, Critically, the RSA ANOVA revealed both a significant main effects of trial type and ROI and a significant 3-way interaction between trial type, age, and ROI. With respect to the main effect of trial type (*F*(1, 52) = 130.34, *p* < .001), the average pattern similarity was higher for Hits and CRs (*M* = 0.22, *SD* = 0.19) than for Hits and FAs (*M* = 0.13, *SD* = 0.10). There was also a main effect of ROI (*F*(1.66, 86.12) = 176.25, *p* < .001), with no main effect of age group *F*(1, 52) = 0.12, *p* = 0.73 (*M*_*old*_ = 0.17, *SD*_*old*_ = 0.15; *M*_*young*_ = .18, *SD*_*young*_ = 0.16). Given the collapsed RSA means (i.e., trial type) are not interpretable, we chose not to evaluate these findings further, but to focus on interactions with trial type and age.

To that end, there was a significant trial type by ROI interaction (*F*(1.44, 74.85) = 95.47, *p* < .001), such that the pattern similarity between Hits and CRs was greater than that of Hits and FAs in the AG (Hit and CR: *M*= 0.18, 95% C.I. [0.17, 0.20]; Hit and FA: *M*= 0.11, 95% C.I. [0.10, 0.12]; *t*(53) = 8.55, *p* < .001), IOC, (Hit and CR: *M*=0.37, 95% C.I. [0.35, 0.40]; Hit and FA: *M*=.19, 95% C.I. [0.18, 0.21]; *t*(53) =9.00, *p* < .001), MOC (Hit and CR: *M*=0.47, 95% C.I. [0.44, 0.49]; Hit and FA: *M*= 0.24, 95% C.I. [0.22, 0.26]; *t*(53) =10.63, *p* < .001), medial SFG (Hit and CR: *M* =0.12, 95% C.I. [0.12, 0.13]; Hit and FA: *M*= 0.08, 95% C.I. [0.08, 0.09]; *t*(53) =7.06, *p* < .001), and MTG (Hit and CR: *M*= 0.13, 95% C.I. [0.12, 0.0.13]; Hit and FA: *M*= 0.09, 95% C.I. [0.08, 0.09]; *t*(53) = 9.07, *p=*.03).

There was also an age by trial type interaction (*F*(1, 52) = 9.15, *p*=.004), as well as a significant 3-way interaction (*F*(1.44, 74.85) = 12.03, *p* < .001). The age by RSA type interaction showed that younger adults exhibited higher neural similarity across Hits and CRs (*M*= 0.24, *SD* = 0.20, 95% C.I. [0.22, 0.25]) than Hits and FAs (*M*= 0.12, *SD*= 0.07, 95% C.I. [0.11, 0.12]; *t*(25) = 9.32, *p* < .001). Older adults similarly exhibited higher similarity across Hits and CRs *(M*= 0.21, *SD*= 0.17, 95% C.I. [0.19, 0.22]) than Hits and FAs (*M*= 0.14, *SD*= 0.12, 95% C.I. [0.13, 0.15]; *t*(27) = 6.54, *p* < .001). Breaking down the foregoing 3-way interaction, results showed that the difference between the two RSA metrics was reduced in older adults compared to younger adults in the IOC (*M*_*OA*_=0.12, *M*_*YA*_=0.25), *t*(52) = 3.41, *p* < .005, MOC (*M*_*OA*_=0.16, *M*_*YA*_=0.30), *t*(52) = 3.62, *p* < .001, and medial SFG (*M*_*OA*_=0.027, *M*_*YA*_=0.052), *t*(52) = 2.36, *p* = .022 (the difference in the medial SFG did not survive Bonferroni correction)^2^. [See figure 3].

**Figure 3.**
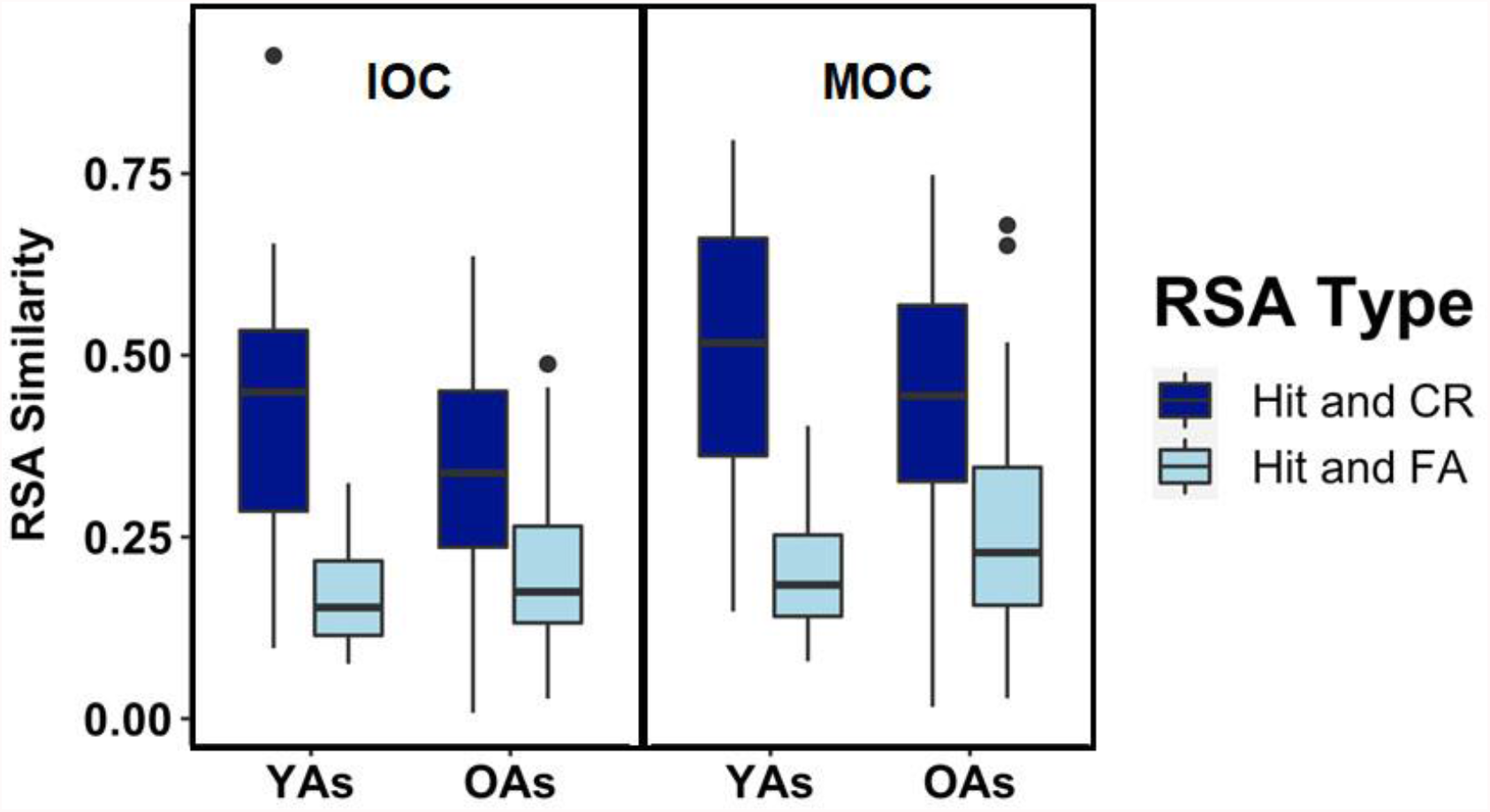
RSA for Hits & False Alarms (FA) and Hits & Correct Rejections (CRs) in Inferior Occipital Cortex (IOC) and Middle Occipital Cortex (MOC) across younger adults (YAs) and older adults (OAs).

Next, we ran a series of linear regressions, including Age, Hit-FA similarity, and the interaction term, to examine how age and Hit and FA similarity may predict false alarm rates. In line with the foregoing behavioral results, age was a significant predictor of false alarm rates in all ROIs such that older adults (*M*=0.29) had higher false alarms than younger adults (*M*=0.16), all *p*’s<.001. In the MOC (*b*=0.36, *p=*.017), MTG (*b*=1.49, *p=*.003), and medial SFG (*b*=2.52, *p<*.001), similarity was a significant predictor of FAs such that as similarity increases, FA rates increase. The age by similarity interaction was not significant in any ROI^3^.

A similar set of linear regressions using Hit-CR similarity in place of Hit-FA similarity, were run within the ROIs to predict corrected recognition rates. Neither Hit-CR similarity nor the interaction term was found to predict corrected recognition rates (all *p’s*>.15). Mirroring the effect of age in the FA regression above, age was a significant predictor of correct rejection rates in all ROIs such that younger adults had higher correct rejections (*M*=0.45) compared to older adults (*M*=0.26), all *p*’s<.001.

## Discussion

The current study aimed to shed light on the neural processes underlying the role of false memories within age-related associative memory deficits. Behaviorally, our results revealed the observed associative memory deficit in *d’* was driven by greater false alarm rates in older adults than younger adults, rather than by an age-related reduction in hit rates. Our ERS analysis reveal a significant main effect of ROI and configural congruency condition, such that we observed an advantage for representing and processing targets as a function of intact configural congruency across memory phases within occipital regions. Yet, absent of age effects in these regions, the results suggest that older adults, like young, exhibited greater correspondence of neural activity between encoding and retrieval for targets that maintained the same configural context across memory phases than targets that did not. With regard to the retrieval-related RSA, we saw an age-related difference in the similarity of neural patterns underlying accurate and inaccurate memory retrieval. Specifically, results revealed that pattern similarity between Hits and FAs and Hits and CRs differed as a function of age in occipital ROIs, such that older adults exhibited a smaller difference between the two similarity metrics than did younger adults. Additionally, pattern similarity between Hits and FAs correlated with FA rates across a number of regions in older adults. Behavioral and multivariate findings will be discussed in turn.

### Behavioral Results

Consistent with a wealth of past research (Chalfonte & Johnson, 1996; Naveh-Benjamin, 2000; Naveh-Benjamin, Guez, Kilb, & Reedy, 2004; Old & Naveh-Benjamin, 2008a; Overman & Becker, 2009; Overman et al., 2018; Rhodes et al., 2008), older adults exhibited deficits in associative memory compared to younger adults. Specifically, older adults showed overall lower behavioral discriminability, measured by *d’* (see Figure 3). When breaking down *d*’ and examining hit and false alarm rates, results showed that the foregoing age reduction in *d’* was driven not by age differences in hit rates, but by age-related increases in associative false alarms. Absent of any interactions with age, results suggest that the foregoing age differences were not affected by either encoding or retrieval conditions but reflect a general decline in the ability to identify novel information during retrieval. While age-related associative memory deficits are often conceptualized with respect to errors in target detection, the current finding is consistent with past research that also finds that errors in lure rejection (i.e., false memories) underlie age-related associative memory deficits (e.g., Castel & Craik, 2003; Healy et al., 2005; Naveh-Benjamin et al., 2003; Old & Naveh-Benjamin, 2008b; Overman & Stephens, 2013; Rhodes et al., 2008).

Moreover, this pattern of results is consistent with the fact that older adults tend to make memory responses based on a feeling of familiarity opposed to recalling specific details during memory retrieval (e.g., Bastin & Van der Linden, 2006; Davidson & Glisky, 2002). In the present memory study, lure familiarity arises due to the fact that the lures were composed of recombined pairs from encoding, such that all *items* within a lure pair are ‘old’, yet not the specific association amongst the items. In order to identify a recombined lure as new, one must overcome the item familiarity and respond based on the novel combination of items (Cohn & Moscovitch, 2007; Lepage, Brodeur, & Bourgouin, 2003; Rotello & Heit, 2000). The current behavioral results suggest that there is an age-related decline in the ability to undertake this relatively difficult task of identifying the novel association as new. This is also consistent with an age-related reduction in strategic processing at retrieval (Cohn, Emrich, & Moscovitch, 2008), as well as with previous work showing that older adults have difficulty employing strategic processing at retrieval to reject similar but novel information (i.e., lack of recall-to-reject processing) (Bowman & Dennis, 2015; Pierce, Waring, Schacter, & Budson, 2008). Future research may look at different response profiles across age based on whether the option of responding ‘rearranged’ is offered (i.e., ‘intact’ vs. ‘rearranged’), as is done in some associative memory tasks.

### ERS analyses

To understand why older adults endorse associative lures as ‘old’ pairs at higher rates than younger adults, we first examined whether there were age differences in how encoded information related to targets was recapitulated during retrieval. Specifically, following earlier work with younger adults (Gerver et al., 2020) we examined the similarity of neural patterns of target pairs as a function of both the encoding configuration and configural congruency across memory phases. We hypothesized that, consistent with prior research in item and source memory (Chamberlain, Bowman, & Dennis, under review; Folville et al., 2020; Hill et al., 2021; Trelle et al., 2019), older adults would exhibit reduced ERS of target pairs compared to younger adults. We theorized that such a reduction in the transfer of associative information from encoding to retrieval would contribute to the reduced distinctiveness of targets and lures within retrieval itself. Our results showed that in older, like younger adults (Gerver et al., 2020), both configural context and configural congruency were critical factors in ERS within the IOC and MOC (see figure 2). Specifically, these findings suggest that when an associative target is presented in the same configural context across both memory phases, it evokes greater recapitulation of visual processes associated with the encoding episode than if the configural context is altered for the item-item pair. This pattern of ERS across target conditions in older adults suggests that, like younger adults (Gerver et al., 2020), older adults take advantage of stability in the configural context across memory phases when reinstating memory representations associated with associative pairs across memory phases. This finding aligns with the behavioral results showing higher hit rates in the congruent compared to incongruent configural condition across age groups. While we didn’t find that that ERS correlated with behavioral metrics of associative memory success or false memory rates, it is likely that this consistency for configurally-congruent pairs supports identification of targets using both item-item details and general familiarity of the associative configurations.

Noted above, our Bayes Factor analysis examining age differences within the two occipital ROIs found that while there is moderate evidence supporting the null hypothesis (no age differences) within the MOC, there is moderate evidence supporting the alternative hypothesis (age differences) within the IOC. A closer look at the ERS values within IOC suggests that this age difference may lie in the fact that, despite exhibiting the same pattern of ERS across the four conditions, older adults exhibit somewhat lower overall ERS for the two incongruent conditions compared to younger adults. Thus, the current results offer mixed evidence for an age-reduction in target reinstatement within the occipital cortex. Prior work in the field of memory and aging has observed age-related decline in the reinstatement of target processing, (Chamberlain et al., under review; Folville et al., 2020; Hill et al., 2021; Trelle et al., 2019) within occipital cortices. This prior memory research concludes that aging is associated with reduced reinstatement of neural activity associated with the encoding of detailed stimuli.

Irrespective of this deficit, prior findings also suggest that this decrease in recapitulation does not affect the subjective vividness of these memories in older adults (Folville et al., 2020) and may be related to neural dedifferentiation during encoding (Hill et al., 2021). Given the complicated nature of age differences within the ERS results, future work should continue to examine how congruency across memory phases affects the reinstatement of associative information in aging.

One important point to be considered with respect to any group comparisons in ERS metrics is that ERS metrics do not account for the specificity of the information that is carried over from encoding to retrieval or whether the same type of processing is occurring within the same ROIs across age groups. They merely account for similarity in neural patterns within an individual across memory phases. Thus, it cannot be concluded that both age groups encode and maintain the same information from encoding to retrieval, simply that both age groups exhibit relatively similar patterns of encoding and retrieval activity across the two memory phases. Thus, it may be that younger adults encode and maintain more nuanced or detailed associative representations compared to older adults, whereas older adults process information in a more generic or gist-based manner (e.g., Pierce, Sullivan, Schacter, & Budson, 2005; Schacter, Koutstaal, & Norman, 1997; Stephens & Overman, 2018; Tun, Wingfield, Rosen, & Blanchard, 1998) and that we cannot capture these subtle differences in with the current ERS analysis approach. With more advanced analyses, we will hopefully be able to assess the specificity of the memory representations across age groups.

### RSA analyses

While the investigation of target-related patterns of activity indicates stability of the target representation across age groups, the investigation of age differences in pattern similarity related to false alarms during retrieval told a different story. Specifically, to investigate the neural processes underlying older adults’ reduced ability to distinguish between targets and lures, we used pattern similarity analyses to determine how the relationship between hits and false memory processing differs from that of hits and correctly rejected lures. We also directly investigated whether this relationship was affected by age and whether neural similarity across targets and lures related to false memory rates within our participants. Results showed that while the neural pattern similarity underlying hits and correct rejections was greater than neural pattern similarity underlying hits and false alarms, this difference between the two similarity scores was reduced for older adults. Specifically, the difference between pattern similarity underlying Hits and CRs and Hits and FAs was reduced in older compared to younger adults (see Figure 3), with overall age differences driven by neural processing within both the IOC and MOC (see Figure 3).

The finding that neural patterns underlying veridical memories, in any form, are more similar than those corresponding to a behavioral response of ‘old’, may be reflective of overlap that exists between target identification and recall-to-reject strategies in recognition memory (Brainerd, Reyna, Wright, & Mojardin, 2003; Gallo, Bell, Beier, & Schacter, 2006; Rotello & Heit, 2000). That is, in order to overcome the familiarity inherent in targets and recombined lures in associative memory retrieval, one needs to recall the specific correspondence between the items within the association and not merely the past occurrence of the individual items. The current results suggest that when this process of lure rejection is successful, individuals, irrespective of age, exhibit higher similarity between the identification of intact target pairs and the identification of the recombined novel pairs as lures than they do with novel lure pairs that are erroneously identified as old.

Critically, with respect to aging, the results also identified a reduction in the difference between the RSA metrics in the IOC and MOC in older compared to younger adults. Driven by an increase in the neural overlap across Hits and FAs, within older adults, this suggests that the two trial types may be less distinct in how they are processed in older adults, specifically within occipital regions. This finding is consistent with a wealth of literature showing that older adults exhibit dedifferentiation with respect to the similarity of neural processing across distinct trial types (e.g., Deng et al., 2021; Koen, Hauck, & Rugg, 2019; Koen & Rugg, 2019; St-Laurent, Abdi, Bondad, & Buchsbaum, 2014). While age-related dedifferentiation is usually reported in the context of distinctiveness of object processing (e.g., face/scenes), our results extend this past work, given the current need to differentiate between two fairly similar sets of objects, specifically, intact and recombined pairs. This finding is also consistent with prior work in our lab indicating that patterns of neural activation between targets and lures are less distinguishable in visual cortices in aging (Bowman et al., 2019). In this past work, age deficits in memory were related to reduced discriminability of cognitive processes (i.e., old/new recognition). To that point, and in line with our predictions, the similarity in neural patterns of activity underlying Hits and FAs predicted FA rates in our sample within several ROIs, including MOC, MTG, and medal PFC (see Figure 4). While activity in the foregoing regions has been linked to false memory processing across a large number of studies in both young and older adults (e.g., Dennis, Bowman, et al., 2014; Dennis, Kim, & Cabeza, 2008; Dennis & Turney, 2018; Duarte, Graham, & Henson, 2010), the current finding supports the idea that similarity in neural processing across Hits and FAs is likely to underlie the fact that lures are erroneously labeled as “old”.

**Figure 4.**
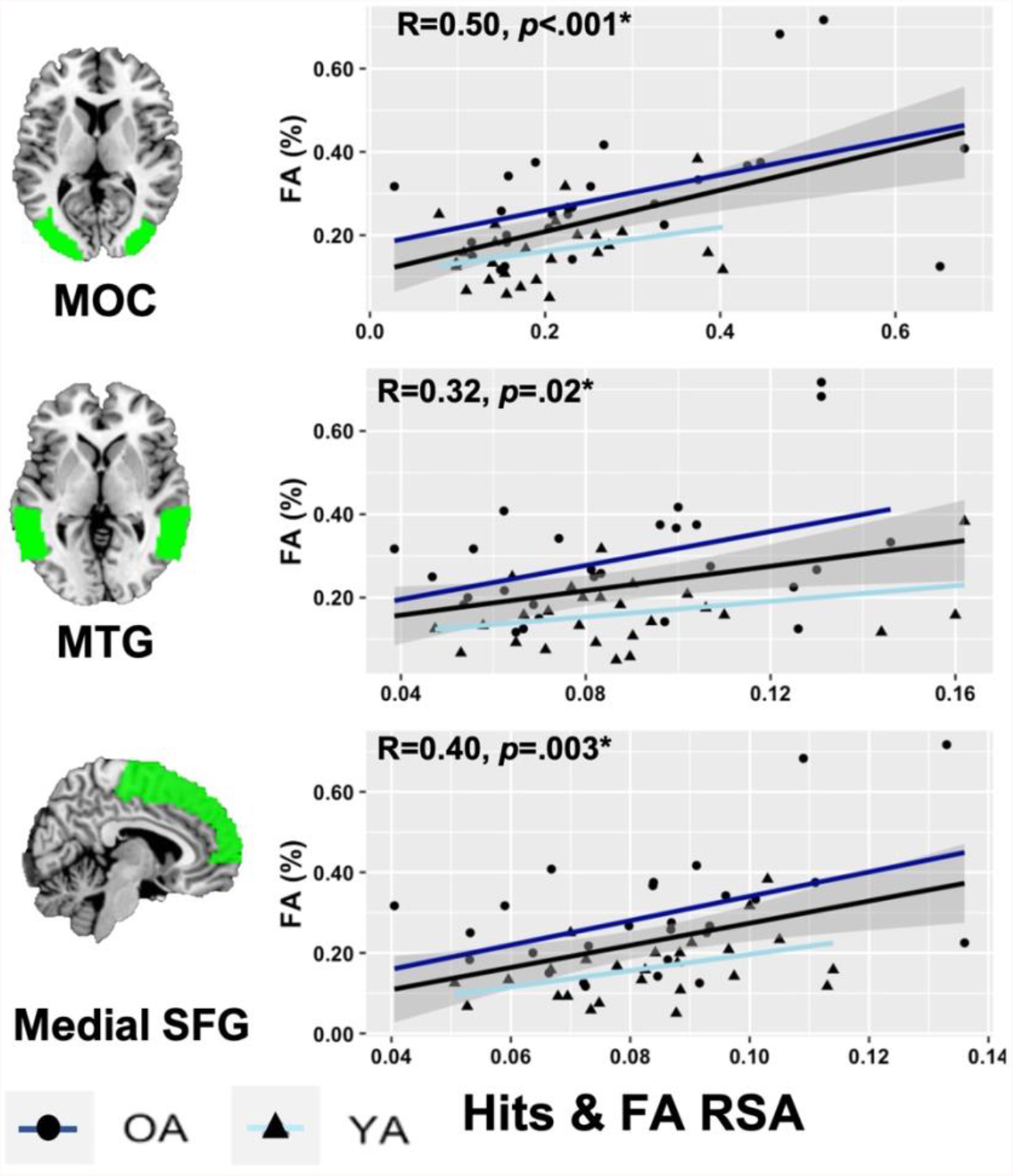
Correlation between RSA for Hits & False Alarms (FA) with FA rates in each ROI collapsed across age (YAs: younger adults; OAs: older adults). Group regression lines are also included for illustrative purposes. Middle Occipital Cortex (MOC); Middle Temporal Gyrus (MTG); Medial Superior Frontal Gyrus (SFG).

This conclusion is consistent with a wealth of past research examining false memories and aging that points to perceptual relatedness, shared semantic gist, and overlapping mnemonic representations across targets and lures as underlying mechanisms accounts for age-related increases in false memory rates (e.g., Dennis, Bowman, et al., 2014; Devitt & Schacter, 2016; Tun et al., 1998). In accord with the factors described above, associative lures, much like related lures within semantic and perceptual false memory paradigms (e.g., DRM test, perceptual relatedness studies; see Dennis, Bowman, & Turney, 2015), share a high degree of similarity with target items, as both items within the rearranged pair have been previously encountered. Thus, the lure incurs a high perceptual overlap with encoded information and is likely to evoke a similar perceptual and conceptual gist as to that of target items. In the absence of clear and distinct representations across trial types, such shared perceptual and gist-related representations are likely to lead to a similar lure being incorrectly labeled as “old”. Regarding episodic representations, the MTG has been shown to be involved in processing the gist associated with episodes (Simons et al., 2005; Wise & Price, 2006) and MOC in processing more general properties of stimuli (Slotnick & Schacter, 2004, 2006; Vaidya, Zhao, Desmond, & Gabrieli, 2002). Given the correspondence in the relationship between Hit and FA representations and false memory rates in these regions, results suggest that the extent to which related lures invoke the general feeling of oldness associated with past veridical item-item associations, the more likely they will evoke an erroneous response of “old” at retrieval.

Yet the similarity of representations within these lower-level processing regions does not solely account for the creation of this memory error. The current results also suggest that similarities in the manner in which a Hit and FA are processed within higher level monitoring and evaluation regions, including both the medial SFG and parietal cortex, may contribute to increased false alarm rates in older adults. The correlation between Hit and FA RSA and false memories in the medial SFG is particularly interesting given that this region has consistently been identified in false memory studies across a wide variety of false memory paradigms (for a metanalysis see Kurkela & Dennis, 2016), including studies involving older adults (e.g., Dennis, Bowman, et al., 2014; Dennis & Turney, 2018; Webb & Dennis, 2019). Noted above, the role of the medial SFG during false memory retrieval has been linked to part of a frontal-parietal cognitive control network responsible for the evaluation and monitoring of critical lures (Atkins & Reuter-Lorenz, 2011; Dennis, Bowman, & Vandekar, 2012; Dennis, Johnson, & Peterson, 2014; Garoff-Eaton, Kensinger, & Schacter, 2007; Hofer et al., 2007; Iidaka, Harada, Kawaguchi, & Sadato, 2012; Kensinger & Schacter, 2006; Slotnick & Schacter, 2004; von Zerssen, Mecklinger, Opitz, & von Cramon, 2001). Prior findings have established a link between processing within the medial prefrontal cortex and top-down retrieval monitoring in the absence of a strong sensory signal (Dennis, Bowman, et al., 2014) in the face of competing representations (Iidaka et al., 2012), and when there is a strong conceptual similarity between targets and lures (Garoff-Eaton et al., 2007). The current results extend this univariate work from younger adults to show not only that activity within this region support false associative memories in aging, but also that the greater the extent to which lure processing within the medial SFG corresponds to that underlying a veridical memory response, the greater the likelihood of a false memory.

## General Conclusions

The current results shed light on the neural processes underlying age-related associative memory deficit. Behaviorally, our results highlight the role of false memories of recombined pairs at retrieval as a significant, contributing factor to age deficits in associative memory performance. That is, the associative memory deficit was driven by greater false alarm rates in older adults, rather than by a reduction in the hit rate. Paralleling the absence of age deficits in hit rates, we also found no age-related differences in the relationship between patterns of neural activity underlying targets across memory phase (as measured by ERS), though we did see an advantage for representing and processing targets as a function of intact configural congruency across memory phases. Specifically, older adults, like young (Gerver et al., 2020), exhibited greater correspondence of neural activity between encoding and retrieval for targets that maintained the same configural context across memory phases than targets that did not. The results highlight the strength of configural arrangements as part of the associative memory trace and the importance of maintaining configural congruency across memory phases as a means to bolster the transfer of associative information across phases in older adults. This finding is consistent with earlier behavioral work from our collaborative group (Overman, Dennis, McCormick-Huhn, Steinsiek, & Cesar, 2019; Overman et al., 2018) where we found that both the manner of encoding presentation as well as congruency of configural display was critical to memory success. Importantly, the current results suggest that the strength of this representation is preserved across the adult lifespan.

At the same time, both age groups exhibited overall lower pattern similarity for hits and FAs when compared to hit and CRs. This representational difference between a consistent behavioral response (i.e., “old”) and a correct memory decision suggests that there is more neural uniformity associated with veridical memory decisions across the associative memory network than for trials that share a similar behavioral response but differ with regard to memory veracity. Despite this overall difference, results also showed this difference to be reduced in older compared to younger adults, specifically withing occipital regions. Together with prior examining dedifferentiation and neural distinctiveness within false memory paradigms, this finding is indicative of an age-related increase in the similarity in processing across targets and lures that underlies lures being erroneously identified as targets in older adults. To this point, it was the overlap in neural pattern similarity across hits and FAs that correlated with false memory rates across several ROIs. This latter finding speaks directly to the idea that, when both young and older adults experience a high correspondence in neural processing between targets and related lures, they are more apt to erroneously identify the lure as previously studied. Thus, the current results continue to underscore the importance of understanding errors of commission in aging in order to account for age-related memory decline.

## Acknowledgements

We thank Kayla McGraw, M. Andrew Rowley, Joanna Salerno, Harini Babu, and Chloe Hultman for help with data collection and analyses. We also thank Jordan Chamberlain, Dan Elbich, and Ashley Steinkrauss for analysis support and help with earlier versions of this work. This work was supported by the National Institutes of Health under grant R15AG052903 awarded to A.A.O. and N.A.D. In addition, N.A.D. was also supported in part by National Science Foundation grants BCS1025709 and BCS2000047. Portions of the research in this article used the Color FERET (Facial Recognition Technology) database of facial images collected under the FERET program, sponsored by the Department of Defense Counterdrug Technology Development Program Office.

## Footnotes

1 The current sample size in in line with several previous studies examining age-related differences in neural similarity during episodic memory (Folville et al., 2020; Hill, King, & Rugg, 2021; Srokova, Hill, Elward, & Rugg, 2021; Trelle, Henson, & Simons, 2019)

2 The RSA values broken down by age group are reported in Supplemental Table A for descriptive purposes.

3 Despite the lack of age by similarity interactions, we report correlations with behavior for each age group in Supplemental Table A for descriptive purposes. We also plot age group regression lines alongside group regression lines displayed in Figure 4 for illustrative purposes.

