## Supplemental Table A for "Understanding associative false memories in aging using multivariate analyses"

| | Hit & CR Mean (SD) | Hit & FA Mean (SD) | *t* value |
|---|---|---|---|
| **IOC^** | | | |
| OA | 0.33 (0.15) | 0.21 (0.12) | 3.31** |
| | r = 0.18 | r = 0.29 | |
| YA | 0.41 (0.18) | 0.17 (0.07) | 6.58*** |
| | r = 0.20 | r = 0.30 | |
| **MOC^** | | | |
| OA | 0.43 (0.17) | 0.27 (0.16) | 3.59*** |
| | r = 0.11 | r = 0.47* | |
| YA | 0.50 (0.19) | 0.20 (0.09) | 7.31*** |
| | r = 0.15 | r = 0.33 | |
| **MTG** | | | |
| OA | 0.11 (0.06) | 0.09 (0.03) | 2.23** |
| | r = -0.003 | r = 0.42* | |
| YA | 0.14 (0.06) | 0.09 (0.03) | 4.01*** |
| | r = -0.10 | r = 0.35 | |
| **HC** | | | |
| OA | 0.05 (0.03) | 0.05 (0.02) | -0.13 |
| | r = 0.14 | r = -0.24 | |
| YA | 0.07 (0.03) | 0.06 (0.01) | 0.63 |
| | r = -0.03 | r = 0.13 | |
| **AG** | | | |

| | | | |
|---|---|---|---|
| OA | 0.19 (0.10) | 0.12 (0.05) | 3.13** |
| | r = 0.01 | r = 0.43* | |
| YA | 0.18 (0.09) | 0.10 (0.04) | 4.04*** |
| | r = -0.12 | r = 0.16 | |
| **Medial SFG^** | | | |
| OA | 0.12 (0.05) | 0.08 (0.02) | 3.62*** |
| | r = -0.02 | r = 0.46* | |
| YA | 0.13 (0.04) | 0.08 (0.02) | 5.10*** |
| | r = -0.21 | r = 0.44* | |
